# C-terminal fusion partner activity contributes to the oncogenic functions of YAP1::TFE3

**DOI:** 10.1101/2025.04.04.647316

**Authors:** Patrick J Cimino, Dylan J Keiser, Abigail G Parrish, Eric C Holland, Frank Szulzewsky

## Abstract

YAP1 gene fusions are found in a multitude of human tumors, are potent oncogenic drivers, and are the likely initiating tumorigenic events in these tumors. We and others have previously shown that a YAP1 fusion proteins exert TEAD-dependent oncogenic YAP1 activity that is resistant to inhibitory Hippo pathway signaling. However, the contributions of the C-terminal fusion partners to the oncogenic functions of YAP1 fusion proteins are understudied. Here, we used the RCAS/tv-a system to express eight different YAP1 gene fusions in vivo and observed significant differences in the latencies of tumors induced by the various YAP1 fusions. We observed that tumors induced by YAP1::TFE3 displayed a significantly different histomorphology compared to tumors induced by other YAP1 fusions or activated non-fusion YAP1. To assess the extent to which the functional TFE3 domains (DNA binding: leucine zipper (LZ) and basic-helix-loop-helix (bHLH); activation domain (AD)) contribute to the oncogenic functions of YAP1::TFE3, we generated several mutant variants and performed functional in vitro and in vivo assays. In vitro, mutation or deletion of the TFE3 DNA binding domains (LZ, bHLH) resulted in reduced TFE3 activity but increased YAP1 activity of YAP1::TFE3. In vivo, deletion of the LZ and bHLH domains did not result in a decrease in tumor incidence but induced the formation of more YAP1-like tumors that lacked prominent features of YAP1::TFE3-driven tumors. By contrast, loss of the TFE3 AD almost completely abrogated tumor formation. Our results suggest that the TFE3 domains significantly contribute to the oncogenic activity of YAP1::TFE3.

## Introduction

YAP1 and its paralog TAZ (encoded by WWTR1) are transcriptional coactivators and potent drivers of cell growth that function through the interaction with several different transcription factors, most prominently TEAD1–4 ^1-5^. The activity of YAP1 is regulated by the Hippo signaling pathway, a cascade of serine/threonine kinases that ultimately phosphorylate YAP1 at several serine residues, resulting in the inhibition of YAP activity. Elevated nuclear YAP1 staining has been observed in several cancers, along with inactivating mutations in upstream Hippo pathway tumor suppressors. Recent cancer genome sequencing studies have identified several gene fusions involving the N-terminal regions of YAP1 or TAZ in different, mainly pediatric, tumor types, including epithelioid hemangioendothelioma (EHE; *WWTR1(TAZ)::CAMTA1, YAP1::TFE3*), supratentorial (ST) ependymoma (*YAP1::MAMLD1, YAP1::FAM118B*), *NF2*-wild type meningioma (*YAP1::MAML2, YAP1::LMO1, YAP1::PYGO1*), fibrosarcoma (*YAP1::KTM2A*), poroma/porocarcinoma (*YAP1::MAML2, YAP1::NUTM1*), thoracic tumors (*YAP1::LEUTX*), and cervical squamous cell carcinoma and endocervical adenocarcinoma (*YAP1::SS18*) ^6-14^. Several of these YAP1 fusions have been detected in multiple different tumor types spread over several anatomical locations and organ sites ^1^, suggesting that both the functionality of the gene fusion and the cell type in which the gene fusion arises can contribute to tumor phenotype and histology.

We and others have recently analyzed the functions of several YAP1 and TAZ fusion proteins and have shown that the expression of these gene fusions in mice is sufficient to induce the formation of tumors ^15-19^. All analyzed YAP1/TAZ gene fusions retain the N-terminal portion of YAP1 and TAZ, respectively, including the TEAD interacting domain (TID) and exert oncogenic YAP1/TAZ activity that is resistant to inhibitory Hippo pathway signaling, as well as necessary and sufficient for the oncogenic functions of these fusions. By contrast, the influence and importance of the C-terminal fusion partners on the functions and oncogenic potential of YAP1/TAZ gene fusions is less understood. This is in part owed to the structural and functional diversity of the different C-terminal fusion partners. While *TFE3, CAMTA1*, and *KMT2A* possess transcription factor activity and contain DNA binding domains ^20-22^, other fusion partners such as *MAML2, MAMLD1, NUTM1*, and *LMO1* (as well as YAP1 and TAZ themselves) lack known DNA binding domains and primarily function as transcriptional co-activators that require the interaction with other transcription factors. In addition to their ability to bind DNA, both *TFE3* and *CAMTA1* (and their respective *YAP1* and *TAZ* fusions) have also been shown to interact with several chromatin remodeling proteins and complexes, such as members of the ATAC complex and EP400 ^18^. However, it remains unknown to what degree the C-terminal fusion partners of YAP1 and TAZ fusions contribute to the aggressiveness and histomorphology of tumors induced by these fusions.

In this study, we intracranially expressed eight different YAP1 gene fusions in vivo to compare their oncogenic abilities. We observed that tumors induced by YAP1::TFE3 significantly differed in their histomorphology from other YAP1 fusion or activated non-fusion YAP1. To study the impact of specific YAP1 and TFE3 domains on the functions of YAP1::TFE3, we generated several point and deletion mutants. In vitro, we observed that deletion of the main YAP1::TFE3 domains (YAP1-TID, leucine zipper (LZ), basic helix-loop-helix (bHLH), and the activation domain (AD)) differentially affected the YAP1 and TFE3 activity of YAP1::TFE3. In vivo, a mutant of YAP1::TFE3 lacking the TFE3 activation domain was completely unable to induce tumor formation. By contrast, mutants harboring reduced TFE3 activity (ΔLZ, ΔbHLH) still showed robust oncogenic activity, although tumors displayed a longer latency and significantly altered tumor histomorphology. Taken together, our results suggest that the TFE3 activity exerted by YAP1::TFE3 significantly contributes to its oncogenic functions.

## Results

### Intracranial expression of different YAP1 gene fusions induces tumors with a variety of histological appearances

We utilized the RCAS/tv-a system for somatic cell gene transfer and cloned *YAP1::TFE3* (YT), *YAP1::MAML2* (YM2), *YAP1::MAMLD1* (YMD1), *YAP1::FAM118B* (YF), *YAP1::NUTM1* (YN), *YAP1::SS18* (YS), *YAP1::PYGO1* (YP), and *YAP1::LMO1* (YL), as well as activated non-fusion NLS-S127/397A-YAP1 (2SA-YAP1) into the RCAS vector (Suppl. Figure S1A-B). Since several of these fusions (YM2, YMD1, YF, YP, YL) are found in central nervous system (CNS) tumors (such as meningioma and ependymoma) ^8,10^, we expressed these YAP1 gene fusions in Nestin-positive cells in the brain of Nestin/tv-a (N/tv-a) *Cdkn2a* null mice. Although YT, YN, and YS have not yet been detected in CNS tumors, several YAP1 fusions (including YM2, YMD1, YF, and YT) have been detected in multiple tumor entities of different organ sites ^1^.

We observed that all analyzed YAP1 fusions as well as 2SA-YAP1 were able to induce tumor formation when expressed in Nestin-positive cells in the brain, albeit at different levels of penetrance (Figure 1A-B; Suppl. Figure S1C). We also observed prominent differences in the histomorphology of tumors induced by the expression of 2SA-YAP1 or the different fusions (Figure 1C; Suppl. Figure S1D).

**Figure 1:**
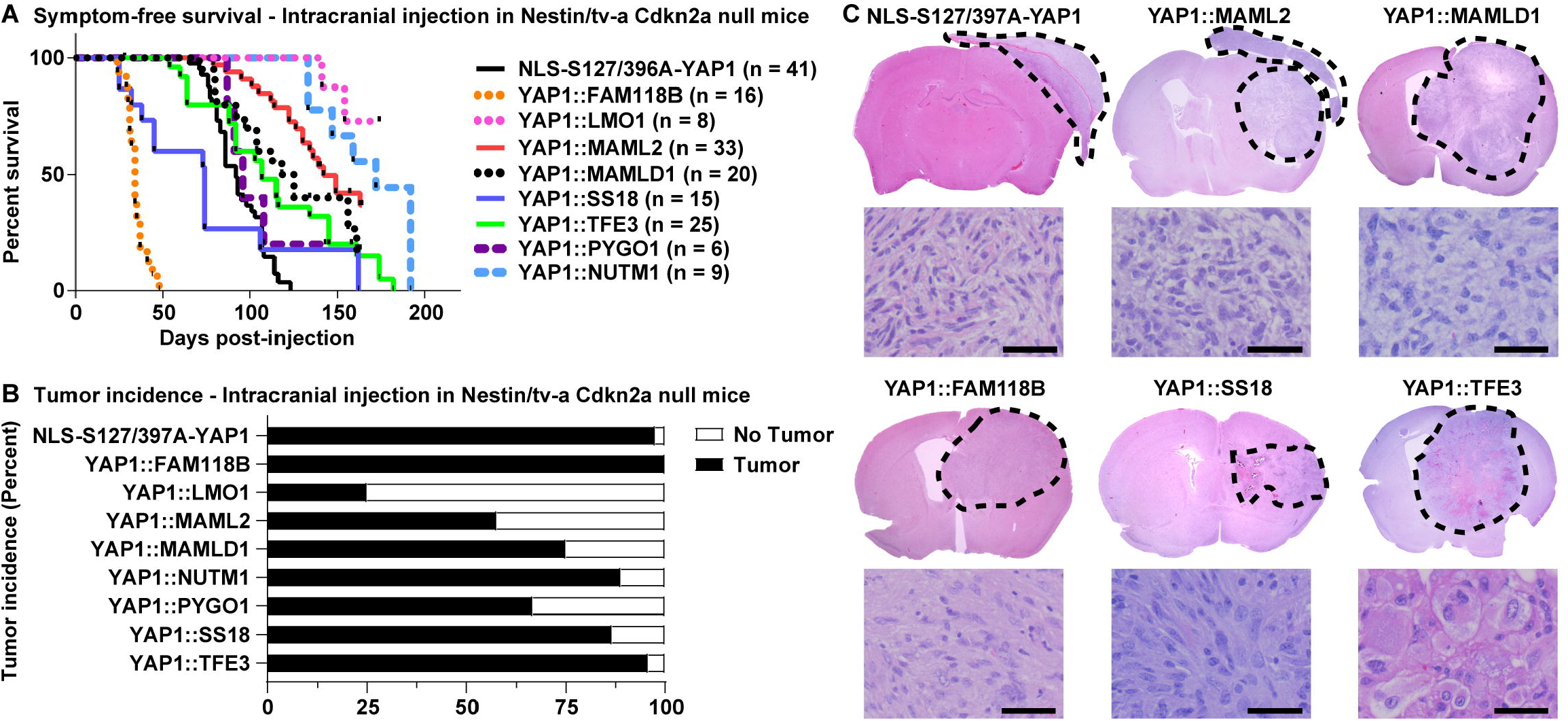
Intracranial expression of different YAP1 gene fusions induces tumors with a variety of histological appearances. A-B) Symptom-free survival (A) and tumor incidence (B) of Nestin/tv-a Cdkn2a null mice intracranially injected to express oncogenic non-fusion YAP1 (2SA-YAP1) or different YAP1 fusions. C) Tumor histology of intracranial tumors induced by the expression of oncogenic non-fusion YAP1 (2SA-YAP1) or different YAP1 fusions. Dotted lines indicate tumor margins. Scale bars indicate 50 μm.

Many of the YAP1 fusion-positive tumors (YMD1, YM2, YS, YF) were large and circumscribed with neoplastic cells that were predominantly spindled-shaped, imparting a sarcomatous-like appearance. Some of these tumors (YMD1, YM2, YF, YS) also had a prominent vasculocentric growth pattern which appeared occasionally as pseudo-infiltrative growth. YF and YS fusion-positive tumors additionally contained patchy areas of tumor necrosis. Tumors induced by YM2, frequently also displayed growth in the extra-axial and extra-cranial planes and had the appearance of meningioangiomatosis-like growth. Mitotic activity was frequently elevated in all *YAP1* fusion-positive tumors. Tumors formed by the non-fusion 2SA-YAP1 had similar spindle cell morphology as the fusion-positive tumors with growth along the convexities in the extra-axial and extra-cranial planes. Rare 2SA-YAP1 tumors showed focal meningioangiomatosis-like growth pattern. Three YAP1 fusions (YN, YP, YL) were unable to effectively induce the formation of intraparenchymal tumors but instead were only able to induce extracranial tumors (Suppl. Figure S1D).

By contrast, YT-expressing tumors displayed a striking difference in histomorphology compared to tumors induced by 2SA-YAP1 or any other YAP1 fusion. YT fusion-positive tumors were intraparenchymal and had a distinct histomorphology. The tumor-brain interface was well-circumscribed (Figure 2A). Patchy areas of lymphocyte aggregates were seen throughout the tumor. The neoplastic cells were strikingly pleomorphic, enlarged and epithelioid, with abundant eosinophilic cytoplasm that demonstrated variable, but frequent lipidized (or xanthomatous-type) changes (Figure 2B-E). Multinucleated cells were also present (Figure 2F). The tumors had high-grade features including elevated mitotic activity, endothelial proliferation, and palisading necrosis. Overall, *YAP1::TFE3* fusion-positive tumors appeared as high-grade gliomas, with histologic features that may be seen in anaplastic pleomorphic xanthoastrocytoma, high-grade glioma with pleomorphic and pseudopapillary features, and epithelioid glioblastoma (Figure 2G-I).

**Figure 2:**
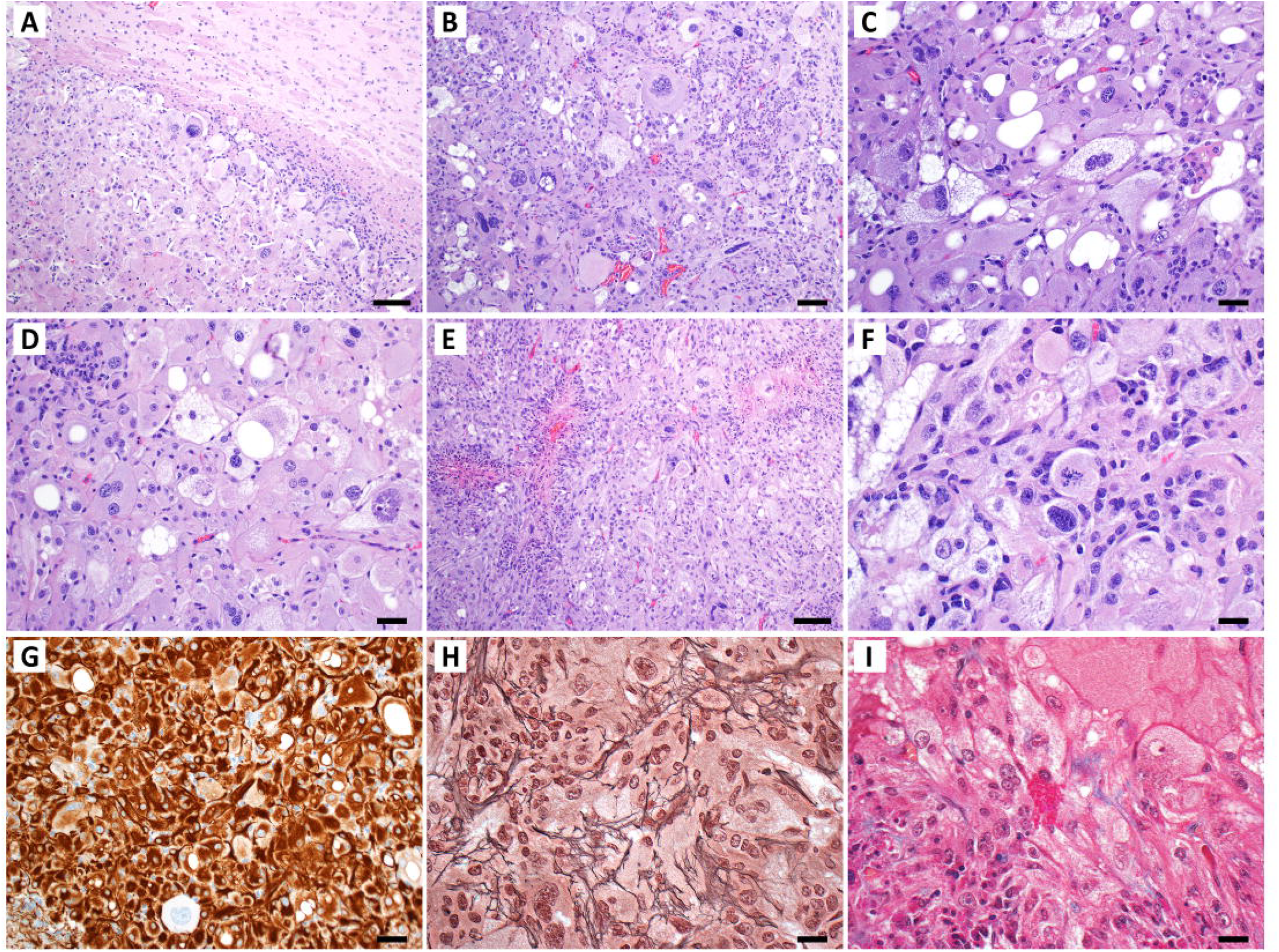
Histopathologic characteristics of YAP1::TFE3 fusion brain tumors. A) Low-power magnification of hematoxylin and eosin (H&E)-stained sections demonstrate that the neoplasm has a well-circumscribed border interfacing with the adjacent brain parenchyma. B-D) H&E-stained sections at higher magnification show the cells are quite pleomorphic with a variable amount of cytoplasmic lipidization. E) For some tumors, there are patchy areas of palisading necrosis. F) Mitotic figures are frequently found. G) A large subset of the neoplastic cells have strong cytoplasmic immunoreactivity for glial fibrillary acidic protein (GFAP). H) Reticulin stain highlights patchy areas of pericellular reticulin fiber deposition. I) Masson’s trichrome stain highlights rare eosinophilic granular bodies (EGBs). Scale bars indicate 20 μm (B-D,F-I) or 50 μm (A,E).

Taken together, these results suggest that the type of YAP1 gene fusion can influence tumor biology and tumors induced by specific YAP1 gene fusions can differ in their histomorphology and aggressiveness.

### TFE3 protein domains influence the YAP and TFE3 activity of YAP1::TFE3

As described above, tumors induced by *YAP1::TFE3* displayed a significantly altered histomorphology compared to tumors induced by the expression of 2SA-YAP1 or other YAP1 fusions.

We have previously shown that YAP1::TFE3 is able to activate both YAP1 and TFE3 target genes and was also able to bind to genomic regions also occupied by wild type TFE3 (wtTFE3), but not by wild type YAP1 (wtYAP1) or other YAP1 fusion proteins ^16^. We decided to study if the activity exerted by the C terminal fusion partner TFE3 contributes to the oncogenic activity of YAP1::TFE3. Several protein domains have been identified within the TFE3 sequence, including a leucine zipper (LZ), basic helix-loop-helix domain (bHLH), and an activation domain (AD) ^23-25^. We generated versions of YT and wtTFE3 harboring either deletions (ΔAD, ΔbHLH, ΔLZ) or mutations (LZmut) in these three domains (Figure 3A; Suppl. Figure 2A). In addition, we generated a version of YT that contained a Ser-94-Ala mutation in the YAP1 TID (S94A-YT) and is unable to bind to TEAD transcription factors.

**Figure 3:**
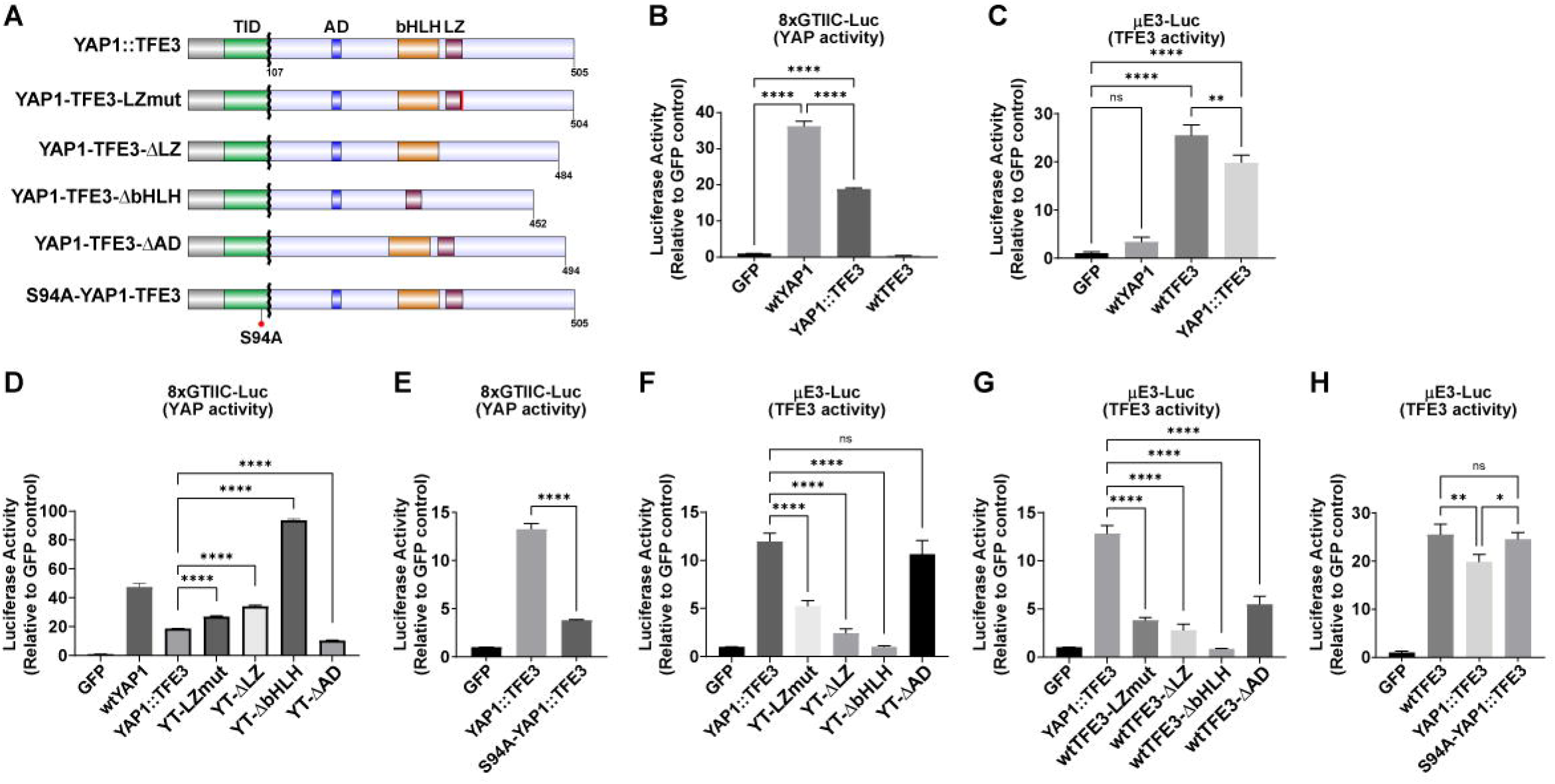
Functional impact of TFE3 domains on the YAP and TFE3-related activity of YAP1::TFE3. A) Schematics of the YAP1::TFE3 sequence and the different truncation mutants. B) YAP1-responsive 8xGTIIC-Luc reporter assay showing the YAP1 activity of wtYAP1, YAP1::TFE3, and wtTFE3 (n = 3). C) TFE3-responsive μE3-Luc reporter assay showing the TFE3 activity of wtYAP1, wtTFE3, and YAP1::TFE3 (n = 3). D) 8xGTIIC-Luc reporter assay showing the YAP activity of the different YAP1::TFE3 deletion mutants (n = 3). E) 8xGTIIC-Luc reporter assay showing the YAP activity of YAP1::TFE3 and S94A-YAP1::TFE3 (n = 3). F-G) μE3-Luc reporter assay showing the TFE3 activity of the different YAP1::TFE3 (F) and wtTFE3 (G) deletion mutants (n = 3). H) μE3-Luc reporter assay showing the TFE3 activity of YAP1::TFE3 and S94A-YAP1::TFE3 (n = 3). Error bars indicate standard deviation. Analysis was done using ordinary one-way ANOVA. (*) *P* < 0.05; (**) *P* < 0.01; (****) *P* < 0.0001.

To assess the baseline activity levels of YT, wtYAP1, and wtTFE3, we performed luciferase assays with the YAP1-responsive 8xGTIIC-Luc reporter as well as the TFE3-responsive μE3-Luc reporter (Figure 3B-C) ^26,27^. Unmutated YT robustly activated both the YAP1-responsive 8xGTIIC-Luc and the TFE3-responsive μE3-Luc reporters, whereas wtYAP1 only activated the 8xGTIIC-Luc but not the μE3-Luc reporter. Conversely, wtTFE3 only activated the μE3-Luc reporter, but not the YAP1-responsive 8xGTIIC-Luc reporter. Of note, YT activated the YAP1-responsive 8xGTIIC-Luc reporter to a significantly lower degree compared to wtYAP1 (18.82 versus 36.2 RFUs, p < 0.0001) and also activated the TFE3-responsive μE3-Luc reporter to a significantly lower degree compared to wtTFE3 (19.82 versus 25.5 RFUs, p < 0.008).

We then assessed the ability of the different YT mutant variants to exert YAP1 activity (Figure 3D-E). Mutation or deletion of the LZ significantly increased the YAP activity of YT (18.67 versus 26.91 and 34.24 RFUs (relative fluorescence units), respectively; p < 0.0001 for both comparisons), whereas deletion of the bHLH domain also dramatically increased the measured YAP activity of YT (18.67 versus 93.7 RFUs; p < 0.0001), almost double the activity of wtYAP1 (47.37 RFUs). By contrast, both the deletion of the TFE3 activation domain (18.67 versus 10.36 RFUs; p < 0.0001) as well as the mutation of the YAP1 TID of YT (S94A-YT; 13.27 versus 3.79 RFUs; p = 0.0009) significantly decreased the YAP activity of YT.

Conversely, we observed that the TFE3 activity of the different YT mutants was generally decreased when disrupting the different TFE3 domains but increased when disrupting the ability of YT to bind to TEADs (Figure 3F-H). Mutation or deletion of the LZ significantly decreased the ability of both YT (11.69 versus 5.24 and 2.43 RFUs, respectively; p < 0.0001 for both comparisons) and wtTFE3 (12.82 versus 3.81 and 2.78 RFUs, respectively; p < 0.0001 for both comparisons) to activate the μE3-Luc reporter. Similarly, deletion of the bHLH domain also dramatically decreased the ability of both YT (11.69 versus 1.03 RFUs; p < 0.0001) and wtTFE3 (12.82 versus 0.85 RFUs; p < 0.0001) to activate the μE3-Luc reporter. Deletion of the TFE3 activation domain significantly reduced the ability of wtTFE3 (12.82 versus 5.49 RFUs; p < 0.0001) but not of YT (11.69 versus 10.63 RFUs; p = 0.12) to exert TFE3 activity. Finally, mutation of the YAP1 TID of YT significantly increased the ability of YT to activate the μE3-Luc reporter (19.82 versus 24.51 RFUs, p = 0.02), resulting in an activity similar to that of wtTFE3 (24.51 versus 25.5 RFUs, p = 0.86).

Taken together, we observed an inverse relationship between the YAP and TFE3 activity exerted by YAP1::TFE3 when deleting several key TFE3 domains responsible for DNA binding. A decrease in TFE3 activity when mutating or deleting the leucine zipper or basic helix-loop-helix domains was generally accompanied by an increase in YAP activity. Likewise, disruption of the ability of YT to bind to TEAD transcription factors resulted in a decreased YAP activity but increased TFE3 activity.

### The TFE3 DNA binding (leucine zipper, basic helix-loop-helix) and activation domains of TFE3 contribute to the oncogenic ability of YAP1::TFE3

We then tested if the different TFE3 domains contribute to the oncogenic tumor forming ability of YAP1::TFE3 by expressing the various YT deletion mutants (ΔLZ, ΔbHLH, ΔAD) in Nestin-positive stem and progenitor cells in the brains of N/tv-a Cdkn2a null mice using the RCAS/tv-a system (Suppl. Figure S2B).

We observed that deletion of either the LZ or bHLH domains did not significantly reduce the penetrance of tumors induced by these YT mutant versions compared to unmutated YT (Figure 4A). However, tumors induced by the YTΔLZ and YTΔbHLH mutants displayed a significantly longer tumor latency (median survival (MS) of 107 versus 191.5 (p = 0.02) and 241 days (p = 0.0008)) (Figure 4B). By contrast, the deletion of the TFE3 activation domain almost completely abolished the ability of YT to induce tumor formation (Figure 4A-B). Only two out of ten mice intracranially expressing the YTΔAD mutant developed small non-symptomatic lesions that were only discovered after euthanizing the mice at the predetermined endpoint (> 200 days post-injection).

**Figure 4:**
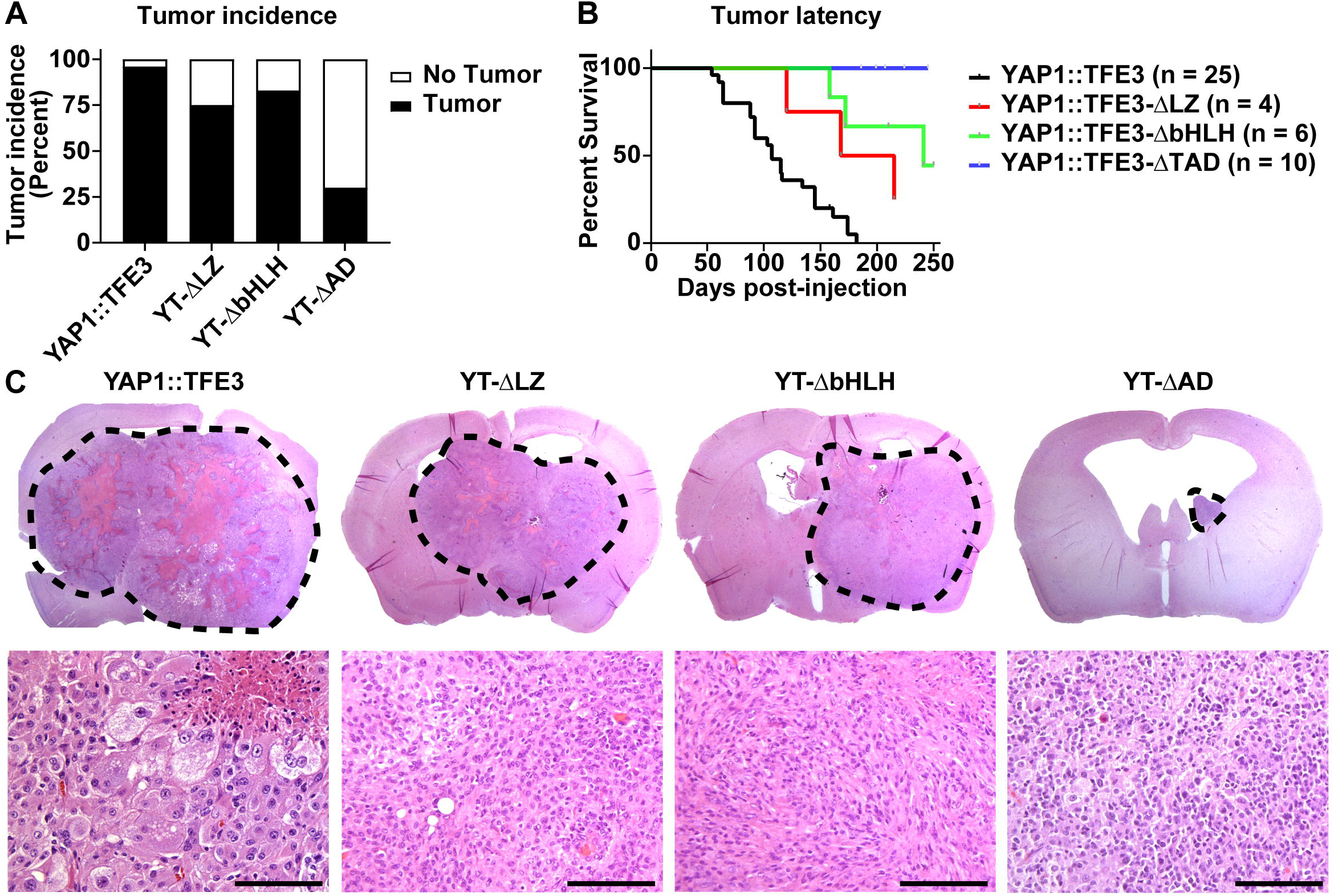
Phenotypical impact of TFE3 domains on the oncogenic functions of YAP1::TFE3. A-B) Tumor incidence (A) and latency (symptom-free survival, B) of tumors induced by the expression of the different YAP1::TFE3 mutants. C) Histomorphological appearance of tumor induced by the expression of the different YAP1::TFE3 mutants. Dotted lines indicate tumor margins. Scale bars indicate 100 μm.

We observed prominent differences in the histomorphology of tumors induced by the expression of the various YT mutants (Figure 4C). As described above, tumors induced by unmutated YT displayed an epithelioid morphology and a high degree of pleomorphism with a large variation in size and shape of tumor cells, resembling the histomorphological appearance of pleomorphic xanthoastrocytomas. Tumors harboring either YTΔbHLH, YTΔLZ, or YTΔAD were also well-circumscribed with high-grade features. However, these *YAP1::TFE3*-modified fusion tumors had neoplastic cells with different cytomorphology. Mainly, they lost their striking pleomorphism and lipidized changes. These neoplastic cells are smaller, more uniform in size and shape, and are epithelioid to spindled.

Taken together, these results suggest that the TFE3 activity exerted by YAP1::TFE3, facilitated by the TFE3 activation domain as well as the LZ and bHLH DNA binding domains, significantly contributes to the oncogenic functions of YAP1::TFE3 and shape the tumor aggressiveness and histomorphology.

## Discussion

YAP1 gene fusions have been identified in a multitude of tumor types spread over a diverse range of anatomical locations. We and others have recently demonstrated that all analyzed YAP1 gene fusions exert oncogenic TEAD-dependent YAP activity that is resistant to inhibitory Hippo signaling ^1,15,16,18^. This deregulated YAP activity itself is necessary and sufficient for the oncogenic functions of YAP1 gene fusions; and the expression of a non-fusion point mutant version of YAP1 (2SA-YAP1) is sufficient to induce the formation of tumors in mice ^16,19^. However, it has become apparent that several YAP1 fusions exert activity that goes beyond that of YAP signaling alone, exemplified by the results of RNA-Seq, Cut & Run/ChIP-Seq, and TurboID experiments that demonstrated additional functions related to the C-terminal fusion partners ^15,16,18,19^. This is further illustrated by the various histomorphologies of tumors induced by the expression of different YAP1 gene fusions as presented by us in this study and previously ^16,19^.

Our results demonstrate that the TFE3 activity of YAP1::TFE3 significantly contributes to the oncogenic functions of YAP1::TFE3. We used an approach in which we deleted or mutated several key TFE3 domains in the YAP1::TFE3 sequence, including the leucine zipper and basic helix-loop-helix domains responsible for the DNA binding abilities of TFE3, as well as the TFE3 activation domain, and subsequently assessed their activity in YAP1 and TFE3-responsive luciferase reporter assays and in vivo experiments. Our results suggest that, although not strictly necessary for the ability of YAP1::TFE3 to induce tumor formation, both the leucine zipper and the basic helix-loop-helix domains significantly contribute to the aggressiveness and histomorphological appearance of YAP1::TFE3-driven tumors. By contrast, the presence of the TFE3 activation domain is essential for the oncogenic functions of YAP1::TFE3, as a mutant version of YAP1::TFE3 lacking the TFE3 activation domain was unable to reliably induce tumor formation. These results are consistent with previous data from our lab on YAP1::MAMLD1, showing that the MAMLD1 transactivation domain is necessary for the activity of YAP1::MAMLD1 ^16^. TFE3 is frequently involved in other, non-YAP1 rearrangements, most commonly found in renal cell carcinoma ^28-30^, and further experimentation will be necessary to assess if our findings on YAP1::TFE3 are translatable to these other TFE3 fusion proteins.

YAP1::TFE3 is primarily found in epithelioid hemangioendothelioma (EHE) as well as clear cell stromal tumor of the lung (CCSTL) and to our knowledge YAP1::TFE3 fusions have not been detected in brain tumors to-date ^12,13,31,32^. Nevertheless, we used the RCAS/tv-a system to express YAP1::TFE3 in neural stem and progenitor cells of the brain to compare it to other YAP1 gene fusions and analyzed the impact of the various functional TFE3 domains on the biology of these tumors. Interestingly, the YAP1::TFE3 CNS mouse tumors we generated had neoplastic cells with dramatic cytoplasmic clearing and vacuolation, which is a histopathologic hallmark of human EHE and CCSTL. We believe that our results will be translatable to the functionality of YAP1::TFE3 in other tumor types.

Future experimentation will be necessary to ascertain how the different TFE3 domains contribute exactly to the functionality of YAP1::TFE3. Previous experiments have demonstrated that both wild type TFE3 and YAP1::TFE3 interact with several protein families, including the ATAC complex ^18^, however, it remains unclear which protein domains exactly facilitate these types of interactions. In addition, our results may have important implications for the future development of inhibitors targeting the functions of YAP1::TFE3. Targeting the leucine zipper or the basic helix-loop-helix domains may not be sufficient to inhibit the growth of YAP1::TFE3-positive tumors, whereas targeting the TFE3 activation domain or the YAP1-TEAD interaction may be more efficacious. We have previously shown that the interaction of YAP1::TFE3 with TEAD transcription factors is essential for its oncogenic functions ^16^.

In summary, our results suggest that the activity exerted by the different TFE3 domains significantly contributes to the oncogenic functions of YAP1::TFE3. These findings enable further insight into how YAP1::TFE3 drives tumor formation and may have important implications for future targeted therapies directed against the YAP1::TFE3 fusion.

## Material and Methods

### Animals

All animal experiments were done in accordance with protocols approved by the Institutional Animal Care and Use Committees of Fred Hutchinson Cancer Center (protocol no. 50842) and followed National Institutes of Health guidelines for animal welfare. Tumors were induced using the RCAS/tv-a system ^33^. Nestin (N)/tv-a;Cdkn2a null mice were used for RCAS-mediated brain tumor formation in this study ^34^. DF1 cells (1 × 10^5^) in a volume of 1 μL were injected near the ventricles into newborn (p0-p2) pup brains ^34^. Mice were monitored until they developed symptoms of disease, such as visible tumors, lethargy, poor grooming, weight loss, dehydration, macrocephaly, seizures, jumping, or paralysis, or until a predetermined study end point.

### H&E staining, immunohistochemistry stainings of FFPE mouse tissues

For routine tumor histomorphology, mouse brains were formalin-fixed and paraffin-embedded, sectioned, and stained with H&E ^34^.

### Plasmid generation

Primers used for plasmid generation are listed in Table S1A. Additional plasmids used in this study are listed in Table S1B. Due to the size limitations of the RCAS vector, truncated versions of YAP1::MAMLD1, YAP1::MAML2, and YAP1::NUTM1 were generated. Truncated versions of YAP1::MAMLD1 and YAP1::MAML2 have been previously described ^16,19^. A truncated version of YAP1::NUTM1 was generated by truncating the C-terminal parts of NUTM1 (see Table S1A).

### Luciferase Assays

HEK293 cells were cultured in DMEM, 10% FBS, 1% Penicillin/Streptomycin. HEK293 cells were seeded into white 96 well plates at 10,000 cells/well the day prior to transfection. Cells were then transfected with the indicated plasmids and a plasmid containing Renilla (100 ng per well for each plasmid) using Lipofectamin 3000 (Thermo Fisher Scientific) according to the manufacturer’s instructions. Luciferase activity was measured 24 hours after transfection using the Dual-Glo Luciferase Assay System (Promega) on a Veritas Microplate Luminometer.

### Western Blots

Cells were cultured, lysed, and processed for western blotting by standard methods. Proteins were resolved by SDS/PAGE (NuPAGE 10% Bis/Tris; LifeTech) according to XCell Sure Lock Mini-Cell guidelines, blocked with 5% milk/TBST and probed with specified antibodies overnight at 4°C in 2% BSA/TBST. After three TBST rinses, species-specific secondary antibodies were added in 2% BSA/TBST. Blots were rinsed three times with TBST before being developed with Amersham ECL Western Blotting Detection Reagents (GE Healthcare). For a list of antibodies used see Table S1C.

### Quantification and statistical analysis

All statistical analyses were conducted using GraphPad Prism 10 (GraphPad software Inc.). To analyze statistical differences between multiple populations’ comparison, ordinary one-way ANOVA was used. p < 0.05 was considered statistically significant.

## Acknowledgments

We thank Anna Spangler, Denis Adair, Linda Lew, and Kelly Grissom for continued administrative assistance and support throughout these experiments. We thank Debra Kumasaka, James Yan, and Zachary Russell for experimental assistance. We thank Elizabeth Jensen and Dolores Covarrubias at the Fred Hutchinson Genomics Core for help with DNA sequencing. Funding for this study was provided by National Institutes of Health grants U54 CA243125 (ECH) and R35 CA253119-01A1 (ECH), National Institutes of Health grant P30 CA015704 (Fred Hutch/University of Washington Cancer Consortium), and University of Utah Department of Neurosurgery and Huntsman Cancer Institute Start-up funds (FS).

## Author contributions

Conceptualization, PJC, ECH, FS; performed experiments, DK, AGP, and FS; data analysis, DK, AGP, PJC, and FS; original manuscript writing, PJC and FS; review and editing, ECH and FS; funding acquisition, ECH and FS; supervision, ECH and FS. All authors read, reviewed, and approved the manuscript.

## Suppl. Figure legends

**Suppl. Figure S1: Intracranial expression of different YAP1 gene fusions induces tumors with a variety of histological appearances**. A) Schematic overview of the structures of the different YAP1 and YAP1 fusion constructs. B) Western Blot of DF1 cells transfected with different RCAS constructs showing expression of YAP1 and YAP1 fusion constructs. Blot was stained with anti-HA antibody, recognizing the N-terminal HA tag present in all constructs. C) Injection summary of different constructs in Nestin/tv-a Cdkn2a null mice. D) Representative high-power H&E images showing tumor histology of intra- and extra-cranial tumors induced by the expression of YAP1::LMO1, YAP1::NUTM1, and YAP1::PYGO1. Scale bars indicate 50 μm.

**Suppl. Figure S2: Functional impact of TFE3 domains on the YAP and TFE3-related activity of YAP1::TFE3**. A) Western Blot of DF1 cells transfected with different RCAS constructs showing expression of YAP1::TFE3 fusion deletion mutant constructs. Blot was stained with anti-HA antibody, recognizing the N-terminal HA tag present in all constructs. B) Injection summary of different YAP1::TFE3 mutant constructs in Nestin/tv-a Cdkn2a null mice.

## Notes

### Competing Interest Statement

The authors have declared no competing interest.

