## Supplementary material for "C-terminal fusion partner activity contributes to the oncogenic functions of YAP1::TFE3": Suppl. Data

Suppl. Figures S1-S2

Suppl. Figure 1

A

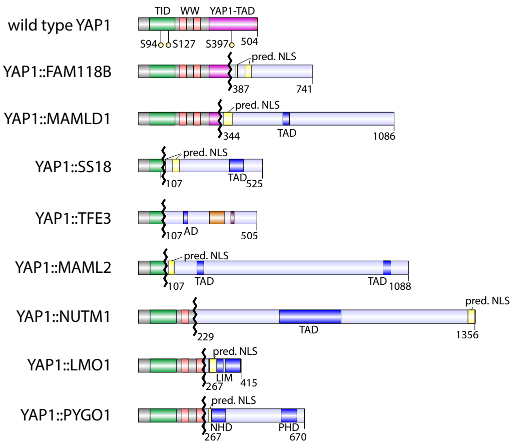


B

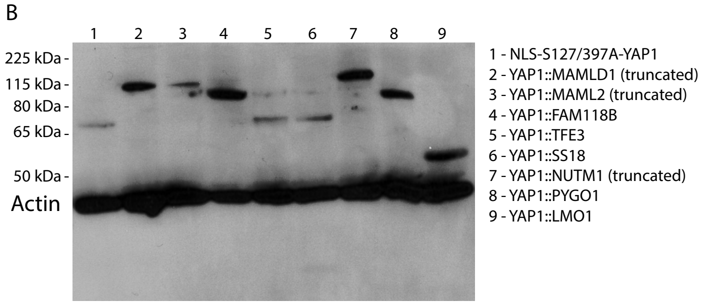


C

| RCAS Vector | No of mice injected | No of mice with tumors | Penetrance (Percent) | Median Survival |
|---|---|---|---|---|
| NLS-S127/397A-YAP1 | 41 | 40 | 97.5 | 92 |
| YAP1::FAM118B | 16 | 16 | 100 | 34 |
| YAP1::LMO1 | 8 | 2 | 25 | n/a |
| YAP1::MAML2 | 33 | 19 | 57.6 | 142 |
| YAP1::MAMLD1 | 20 | 15 | 75 | 121.5 |
| YAP1::NUTM1 | 9 | 8 | 88.9 | 172 |
| YAP1::PYGO1 | 6 | 4 | 66.7 | 96 |
| YAP1::SS18 | 15 | 13 | 86.7 | 74 |
| YAP1::TFE3 | 25 | 24 | 96 | 107 |

D

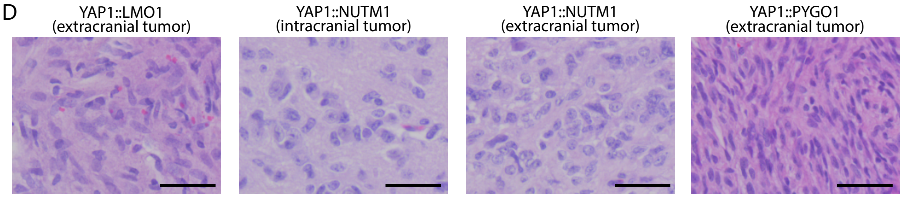

### Suppl. Figure 2

A

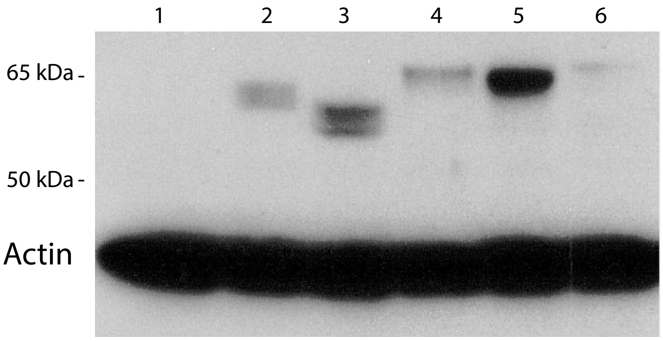


1 - GFP
2 - YAP1::TFE3-ΔLZ
3 - YAP1::TFE3-ΔbHLH
4 - S94A-YAP1::TFE3
5 - YAP1::TFE3-ΔAD
6 - YAP1::TFE3

B

| RCAS Vector | No of mice injected | No of mice with tumors | Penetrance (Percent) | Median Survival |
|---|---|---|---|---|
| YAP1::TFE3 | 25 | 24 | 96 | 107 |
| YAP1::TFE3-ΔbHLH | 6 | 5 | 83.3 | 241 |
| YAP1::TFE3-ΔLZ | 4 | 3 | 75 | 191.5 |
| YAP1::TFE3-ΔAD | 10 | 2 | 20 | n/a |
